# Inter-Individual Variability in Cycling Biomechanics During Experimentally Induced Movement-Evoked Knee Pain

**DOI:** 10.64898/2026.10.09.757760

**Authors:** Sierra Sweeney, Amirreza Hosseini, Shahram Rasoulian, Reza Ahmadi, Koren E. Roach, Junho Park, Amin Komeili, Saied Jalal Aboodarda

## Abstract

Knee pain can alter lower-limb loading and contribute to inter-limb asymmetry, affecting movement mechanics and rehabilitation. Although cycling is frequently prescribed during rehabilitation, the effects of pain on bilateral cycling biomechanics remain poorly understood. This study investigated experimentally induced movement-evoked knee pain on bilateral torque and power output, inter-limb asymmetry, and inter-individual motor responses during cycling. Twenty-three healthy adults (14M and 9F) completed an incremental cycling test followed by eight randomized constant-load trials under pain and no-pain conditions at two intensities. A custom closed-loop torque-dependent electrical stimulation system induced movement-evoked knee pain during the downstroke while bilateral biomechanics were recorded. At higher intensities, movement-evoked knee pain reduced torque (p=0.026, d=0.566) and power (p=0.010, d=0.657) production in the painful limb and altered the normalized symmetry index (NSI) for power (p=0.005, d=0.659), and torque (p<0.001, d=0.754). At lower intensity, pain-leg power (p=0.154) and torque (p=0.104) did not differ significantly between conditions, although torque NSI indicated greater reliance on the non-pain leg (p=0.017, d=0.538). Despite these group-level responses, substantial inter-individual variability was observed ranging from pronounced unloading to increased pain-leg contribution. These findings demonstrate that movement-evoked knee pain alters cycling mechanical output toward the non-painful limb, with substantial heterogeneity in individual motor responses.

## Introduction

Pain is an unpleasant sensory and emotional experience that is linked to, or resembling, experiences associated with actual or potential tissue damage (Raja, et al., 2021). Knee pain is common in musculoskeletal conditions such as knee osteoarthritis, patellofemoral pain syndrome, and ligament or meniscal injuries and is frequently accompanied by altered lower-limb loading and inter-limb asymmetry (Parkinson, Apps, Morris, Barnett, & Lewis, 2021). Although minor asymmetries of less than 10% may occur in healthy individuals, larger discrepancies can be observed in clinical populations. Such asymmetries may alter movement mechanics and contribute to compensatory loading of the contralateral limb (Gustafson, Anderton, Sowa, Piva, & Farrokhi, 2018; Metcalfe, et al., 2012; Hummer, Porter, Cates, & Zhang, 2024; Buddhadev, Crisafulli, Suprak, & Juan, 2018). Cycling is frequently prescribed as a low-impact rehabilitation exercise for lower-limb musculoskeletal conditions; however, joint pain may alter bilateral force production during cycling, leading to compensatory movement patterns. While previous studies have sought to characterize the effect of pain on movement biomechanics, the isolated effects of knee pain on bilateral cycling mechanics remain poorly understood. Studying these effects is challenging, primarily because pain is inherently variable and subjective (Fillingim, 2017), and often accompanied by confounding factors, such as long-term pain-alleviating neurophysiological adaptations (Koop, Schipholt, Scholten-Peeters, & Coppieters, 2021).

To explore the effect of isolated lower limb pain on human movement responses, previous investigators have used different experimental pain models, including muscle-centric pain interventions, such as blood flow restriction (Azevedo, et al.; Zhang, Abel, Macphail, & Aboodarda; Zhang, et al., 2025; McClean, Zhang, Khaledi, Yacoud, & Aboodarda, 2023), or hypertonic saline or capsaicin injections into active muscles (Smith, Micklewright, Winter, & Mauger, 2020; O’Malley, Smith, Mauger, & Norbury, 2024; Norbury, Smith, Burnley, Judge, & Mauger, 2022). Although these approaches can reliably induce experimental muscle pain, the nociceptive stimulus originates primarily from muscle tissue rather than the knee joint. Therefore, biomechanical responses observed with these models may not reflect specific effects of knee joint pain on lower-limb mechanics. Electrical stimulation applied around the patellar region has consequently been used to induce localized and transient knee pain without producing tissue damage. Importantly, this experimental pain model allows for controlled manipulation of pain intensity and timing during movements (Cabral, et al., 2023). Using this model during walking, Charlton et al. (Charlton, et al., 2024) reported compensatory changes in locomotor biomechanics, while Gallina et al. (Gallina, et al., 2024) demonstrated alterations in motor responses and muscular coordination. Broadly, the adaptive responses observed across these studies were interpreted as protective motor strategies adopted by individuals to reduce pain while preserving task performance in the short term (Merkle, Sluka, & Frey-Law, 2018).

The magnitude and temporal pattern of perceived pain may influence the extent and nature of these protective motor adaptations, with consequent effects on joint loading and motor output variability. Movement-evoked pain refers to pain that is provoked or aggravated by movement and may vary with the mechanical demands of the task (Cabral, et al., 2023). Cabral et al. (Cabral, et al., 2023) developed a force-dependent experimental pain model in which stimulation intensity increased with knee-extension torque during submaximal isometric contractions. Movement-evoked knee pain produced substantial reductions in force production, demonstrating that coupling pain intensity to mechanical output can provide a controlled method for examining pain-related motor adaptation. However, this work was limited to a single-joint isometric task. Extending a torque-dependent movement-evoked pain model during cycling minimizes confounding variables associated with pathological conditions and allows direct assessment of pain-related adaptations during a dynamic, bilateral, multi-joint task that is directly relevant to rehabilitation.

Another important consideration is that motor responses to pain are not uniform across individuals. Previous research has demonstrated substantial inter-individual variability in pain-related movement responses, with individuals adopting different strategies to maintain task performance while responding to pain (Merkle, Sluka, & Frey-Law, 2018). Nevertheless, experimental pain studies are commonly interpreted primarily through group-level averages, despite awareness that motor responses to pain are highly heterogeneous (Charlton, et al., 2024; Dionne, Bartoshuk, Mogil, & Witter, 2005; Cabral, et al., 2023; Ducas, Marineau, & Abboud, 2024). To address this limitation, individual-based frameworks have been developed to complement group-based analyses and better characterize the range and direction of responses within a population (Atkinson, Williamson, & Batterham, 2019). Importantly, such approaches should describe response patterns as a continuum rather than impose rigid classifications of “responders” and “non-responders”.

To address these gaps in knowledge, the primary objective of the current study was to explore the effects of movement-evoked experimental knee pain on bilateral torque and power production and inter-limb asymmetry during cycling tasks. It was hypothesized that movement-evoked knee pain would reduce power and torque production in the pain leg, resulting in greater inter-limb asymmetry during cycling. The secondary objective was to characterize inter-individual variability in response to movement-evoked experimental knee pain during cycling tasks. Considering previous evidence demonstrating substantial inter-individual variability in pain-related motor responses, it was hypothesized that the magnitude and pattern of biomechanical responses would vary between individuals. Understanding how pain influences individualized movement responses during cycling is imperative for optimizing rehabilitation applications.

## Methods

### Participants

An a priori sample-size calculation was performed using G*Power (version 3.1). Based on the time-to-peak-torque results reported by Cabral et al. (Cabral, et al., 2023), a within-subjects ANOVA with three conditions, an effect size: 0.50, power: 0.8 and α: 0.05, indicated that a minimum of 9 participants was required. To support the secondary objective of characterizing inter-individual variability in biomechanical responses, 23 healthy, recreationally active adults participated in the study (9F, 14M; age: 24.9 (7.4) years; height: 175.1 (8.1) cm; body mass: 72.1 (10.9) kg).

Participants reported no lower-limb injury within the previous six months and no neurological or musculoskeletal conditions that could influence cycling performance. Before data collection, all participants completed the Physical Activity Readiness Questionnaire Plus (Canadian Society for Exercise Physiology) (Warburton, Jamnik, Bredin, & Gledhill, 2011). The study was approved by the Conjoint Health Research Ethics Board at the University of Calgary (REB23-1111) and conformed to the latest version of the Declaration of Helsinki.

### Experimental Setup

All cycling tasks were performed on a custom, sensor-integrated Echelon EX-5 stationary bike (Echelon Fitness Multimedia LLC, Chattanooga, TN, USA). The system enabled real-time measurements of three-dimensional pedal forces for detailed analysis of bilateral cycling mechanics (Sweeney, et al., 2026). Each pedal integrates a Sensix ICS-MB six-axis force-torque sensor employing six strain gauges (Wheatstone bridges) to measure 3D forces (Fx, Fy, Fz) and moments (Mx, My, Mz). The pedal body (84 mm × 45.2 mm, 450 g) houses a sensor core with measurement ranges of ±1200 N (Fx, Fy), ±4700 N (Fz), and ±90 N.m (Mx, My), ±40 N.m (Mz), with overload tolerances of up to 6470 N for force and 230 N.m for moment.

### Data Acquisition and Signal Conditioning

A dedicated data-acquisition (DAQ) system (Sensix, Poitiers, France) was used for signal collection and conditioning. Pedal signals were routed through a Sensix DAQ populated with six INA2128 instrumentation amplifiers (Texas Instruments, Dallas, TX, USA), using 4.5 V excitation (±0.002 V), a gain of 1370, and analog low-pass filtering at 2000 Hz.

### Electrical Stimulation Setup

A custom Arduino-based closed-loop control system was developed to to achieve precise, repeatable, and dynamically modulated pain stimuli during pedaling. The system interfaced with a constant-current stimulator (Digitimer DS5 Isolated Bipolar Constant Current Stimulator) [17] and continuously modulated the stimulation intensity to the participant’s instantaneous pedal torque.

To minimize the potential influence of limb dominance on pain-related changes in cycling mechanics, electrical stimulation was applied to the right knee in 12 participants and to the left knee in 11 participants. All participants were right-leg dominant, defined as the leg preferentially used to kick a ball (Carcia, Cacolice, & McGeary, 2019). Surface electrodes were positioned over the patellar region (targeting the infrapatellar fat pad, manually identified by palpation). Stimulation was delivered as a continuous 50 Hz sinusoidal waveform, applied exclusively during the downstroke phase of pedaling (0°–180° crank angle).

The hardware architecture consisted of an Arduino UNO R4 WiFi microcontroller interfaced with a 12-bit digital-to-analog converter (DAC; LTC1257), signal-conditioning circuitry (LM6172), and the DS5 stimulator. These components were integrated into a custom printed circuit board (PCB) to ensure signal stability and minimize electrical interference. Real-time biomechanical data, including crank angle and crank torque, were continuously acquired from an instrumented cycle ergometer and processed using a custom Python-based control interface. The normalized stimulation signal was proportional to the instantaneous crank torque, such that stimulation intensity increased as torque increased during the downstroke phase and was discontinued during the unloaded upstroke. The resulting control signal was transmitted to the microcontroller and converted into an analog voltage through the DAC and scaled and shifted by the signal-conditioning circuit to the bipolar voltage range required by the DS5 stimulator. The DS5 generated and applied the corresponding electrical stimulation to the kneecap.

A 50 Hz sinusoidal waveform was selected to provide smooth transitions in stimulation intensity and sustained nociceptive afferent activation while minimizing muscle twitch or startle responses often associated with square-wave pulses. Stimulation intensity was individually calibrated using a verbal numerical rating scale (NRS), where 0 represented no-pain and 10 represented the worst imaginable pain. During calibration, the maximum torque parameter governing the stimulation-voltage output was adjusted to achieve the predefined target pain intensities of 1–2/10 for the low-intensity condition and 5–6/10 for the high-intensity condition. An exponential smoothing filter was applied to limit abrupt fluctuations in stimulation intensity during cycling. A Python graphical user interface (GUI) enabled real-time adjustment of stimulation parameters, live visualization of the waveform, automated data logging for post-hoc analysis, and autozero calibration of the ergometer sensors prior to testing.

### Experimental Protocol

Participants attended two laboratory sessions separated by at least 48 hours. The first session consisted of familiarization with the cycle ergometer and electrical stimulation procedures, followed by an incremental cycling test used to determine peak power output (PPO). After a 3-minute warm-up at 40 Watts (W) and 60 revolutions per minute (rpm), participants completed the incremental test, during which power output increased by 20 W every minute (Ahmadi, Aboodarda, Uchida, Herzog, & Komeili, 2026). The test continued until volitional task failure or until the participant was unable to maintain the required power output for 10 seconds, despite verbal encouragement. The PPO obtained during this session was used to determine individualized workloads for the experimental session. Electrical-stimulation settings determined during the Session 1 were not carried forward to Session 2; stimulation was recalibrated on the experimental day.

The second session involved multiple bouts of constant-load cycling trials under experimental pain (EXP) and no-pain control (CNTRL) conditions. Prior to stimulation, participants were seated still on the bike, and their feet were clipped into the pedals. Based on extensive pilot testing conducted in our laboratory, we found that eliciting a continuum of movement-evoked pain responses during cycling required individualized calibration of stimulation voltage. The stimulation voltage (0-5 V) was scaled between minimum– and maximum-torque parameters in the Python GUI. The minimum-torque parameter was determined while the foot rested on the pedal with the crank in the horizontal position. Stimulation was not delivered below this threshold, thereby preventing stimulation during the upstroke. Participants then cycled at 20% PPO at 60 rpm. The maximum-torque variable was progressively adjusted until a pain intensity of 1–2/10 was reported on a verbal numeric rating scale (0=no-pain, 10=worst imaginable pain). The cycling resistance was then progressively increased by 20% PPO until a pain intensity of 5–6/10 was reported. The workload at which this target pain intensity was reached was recorded as the participant-specific high-intensity workload and was subsequently used for both the HI_EXP and HI_CNTRL trials. The minimum– and maximum-torque and resistance settings were then fixed for the remainder of the session to maintain consistent torque-dependent stimulation.

Participants completed eight randomized 2-minute constant-load cycling trials comprising four experimental conditions: (i) LI_CNTRL: lower-intensity cycling at 20% PPO without stimulation; (ii) LI_EXP: lower-intensity cycling at 20% PPO with stimulation calibrated to evoke a pain intensity of 1–2/10; (iii) HI_CNTL: higher-intensity cycling without stimulation; (iv) HI_EXP: higher-intensity cycling with stimulation calibrated to evoke a pain intensity of 5–6/10. Each condition was performed twice. Trials were performed at a cadence of 60 rpm, with a three-minute recovery between trials. A schematic overview of the protocol is presented in Figure 1.

**Figure 1.**
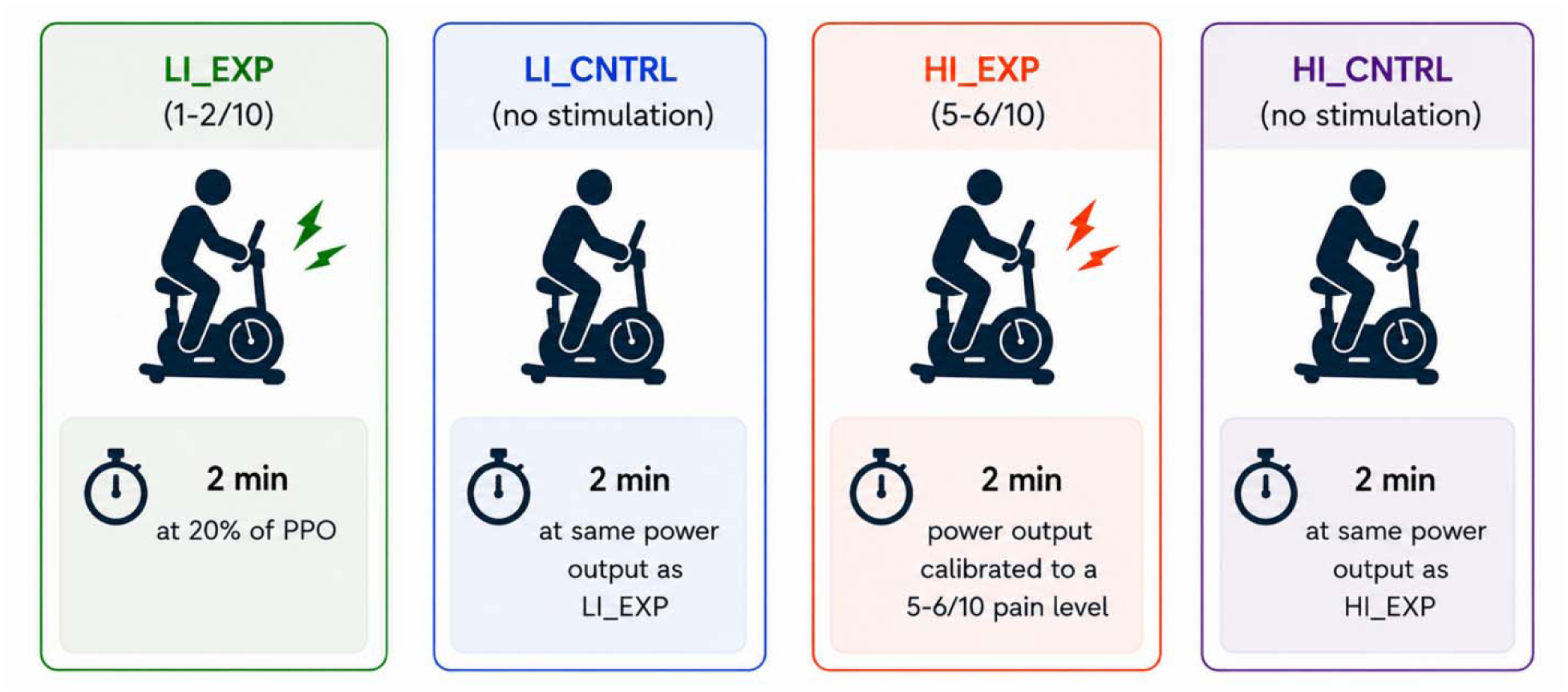
Experimental conditions of cycling trials. HI_CNTRL: higher intensity control condition; HI_EXP: higher intensity experimental condition; LI_CNTRL: lower intensity control condition; LI_EXP: lower intensity experimental condition; PPO: peak power output.

### Data Processing

Torque and power were extracted for each pedal cycle from each leg at five-degree increments of crank rotation. Mean crank angular velocity was calculated from the cycle ergometer. Because electrical stimulation was applied exclusively during the downstroke phase (0–180° crank angle) to avoid cancellation effects associated with averaging across the full pedal cycle. Mean torque and power were therefore calculated across the downstroke phase, where 0° corresponds to top dead center and 180° corresponds to bottom dead center. Mean handlebar force on the pain and non-pain sides and mediolateral seat-post force were extracted over the entire duration of each trial.

### Data Analysis

Inter-limb symmetry was assessed using a normalized symmetry index (NSI), where negative values indicated greater reliance on the non-pain leg and positive values indicated greater reliance on the pain leg. The NSI was calculated separately for torque and power using Eq. 1, where NPL represents the non-pain leg and PL represents the pain leg:

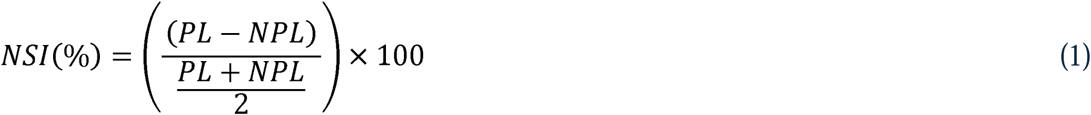

Variability in cycling output was quantified using the coefficient of variation (CV; Eq. 2):

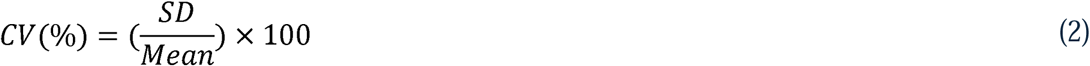

To characterize inter-individual classification of power and torque output, percentage changes between the EXP and CNTRL conditions were calculated using Eq (3).

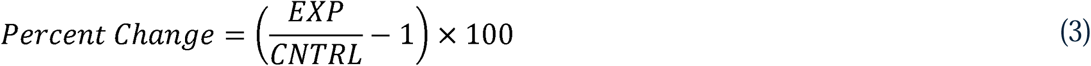

In addition, changes in inter-limb symmetry between conditions were expressed as *ΔNSI* using Eq. 4.

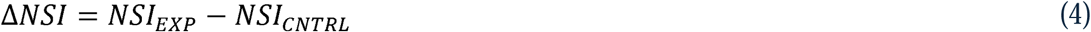

Negative ΔNSI values represented redistribution toward the non-pain leg during EXP conditions, whereas positive ΔNSI values represented redistribution toward the pain leg. This provided a single ΔNSI value for each participant, which was needed to classify individual responses.

Condition-related changes in variability were expressed as the percent change in CV from CNTRL to EXP conditions (ΔCV%; Eq. 5).

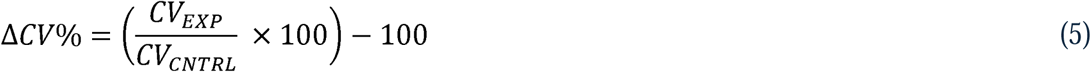

Positive ΔCV% values indicated greater variability during EXP relative to CNTRL conditions, whereas negative ΔCV% values indicated reduced variability.

### Statistical Analysis

Statistical analyses were performed using SPSS Statistics (version 25.0; IBM Corp., Armonk, NY, USA). Data normality was assessed using Shapiro–Wilk tests. Torque, power, CV, handlebar force, and seat post force data satisfied normality assumptions and were therefore analyzed using parametric statistics. However, non-normally distributed variables (NSI) were analyzed using non-parametric tests. Statistical significance was set at *p* < 0.05. The difference in stimulation voltage between lower– and higher-intensity conditions were assessed in 14 participants for whom these data were available using a paired-samples t-test.

To evaluate consistency of torque and power production between the two cycling trials, separate Wilcoxon signed-rank tests were conducted for the pain and non-pain legs at both intensity levels. In addition, intraclass correlation coefficients (ICC2,1) were calculated for each leg, condition (EXP and CNTRL), and intensity (lower and higher) to quantify absolute agreement between repeated trials. Following assessment of between trial differences and reproducibility, the two trials for each condition were averaged for subsequent analyses.

For the variables including pedal torque, power, and CV, separate Bonferroni-corrected two-way repeated-measures ANOVAs were performed with conditions (EXP vs CNTRL) and leg (pain vs non-pain) as within-subject factors. Because NSI values of EXP and CNTRL conditions were not normally distributed, Wilcoxon signed-rank tests were used separately for lower– and higher-intensity cycling.

Handlebar force was analyzed using separate Bonferroni-corrected two-way repeated-measures ANOVA with condition (EXP vs CNTRL) and side (pain vs non-pain) as within-subject factors. Seat post force was analyzed using paired-samples t-tests comparing EXP and CNTRL conditions separately for lower– and higher-intensity cycling. Mean crank angular velocity was compared between CNTRL and EXP conditions using a paired-samples t-test.

When significant interactions were identified, post hoc comparisons were performed using Holm–Bonferroni-corrected paired-samples t-tests. Partial eta squared (ηp²) and Cohen’s d were calculated as measures of effect size for omnibus and pairwise comparisons, respectively (Cohen, 1988).

To characterize inter-individual variability, participants were classified using a distribution-based thresholds derived from the group mean ± 1 standard deviation (SD) of percent-change scores for torque, power, ΔNSI, and ΔCV%. More specifically, these categories were used descriptively to characterize the distribution of responses within the sample and should not be interpreted as a formal identification of “responders” and “non-responders”, consistent with the recommendations of Atkinson et al. (Atkinson, Williamson, & Batterham, 2019). For torque and power, only pain-leg changes below the group mean −1 SD were classified as large decreases in pain-leg output, values between the mean −1 SD and mean as moderate decreases, values between the mean and 0 as small decreases, and positive values as increases.

For ΔCV%, negative values represented reduced variability and positive values represented increases variability during EXP relative to CNTRL. Values below the mean −1 SD were classified as large reductions in variability, values between the mean and mean −1 SD as moderate reductions, values between 0 and the mean as minimal change, values between the mean and mean +1 SD as moderate increases, and values above the mean +1 SD as large increases in variability.

## Results

Mean maximum power achieved on the incremental test was 210.8 (55.1) W (mean (SD)). Lower intensity average power (20% of PPO) was 42.9 (11.0) W. Higher intensity trials, at pain level 5-6/10, were on average 45.0 (9.6) % of PPO at a power level of 95.0 (30.8) W.

ICC analysis demonstrated high reproducibility between repeated trials for torque and power across the pain and non-pain legs and EXP and CNTRL conditions (all ICCs > 0.952; Tables A.1 and A.2). Wilcoxon signed-rank tests identified significant between-trial differences for pain-leg power during the low-intensity CNTRL condition (p=0.036) and for torque and power for the non-pain leg during the high-intensity CNTRL condition (p=0.003 and 0.023 respectively). No other significant between-trial differences were observed. Despite these isolated differences, the high ICC values indicated strong trial-to-trial reproducibility; therefore, the two trials were averaged for all subsequent analyses.

### Mean Crank Angular Velocity

No significant differences were found between CNTRL and EXP conditions for mean crank angular velocity at both low (p=0.401, d=-0.180) and higher intensity (p=0.484, d=-0.150) cycling tasks (Table A.3 in supplemental material).

### Change in Stimulation

A significant difference was observed in voltage supplied to the kneecap at lower as compared to higher intensities (p<0.001, d=1.24) (Figure 2).

**Figure 2.**
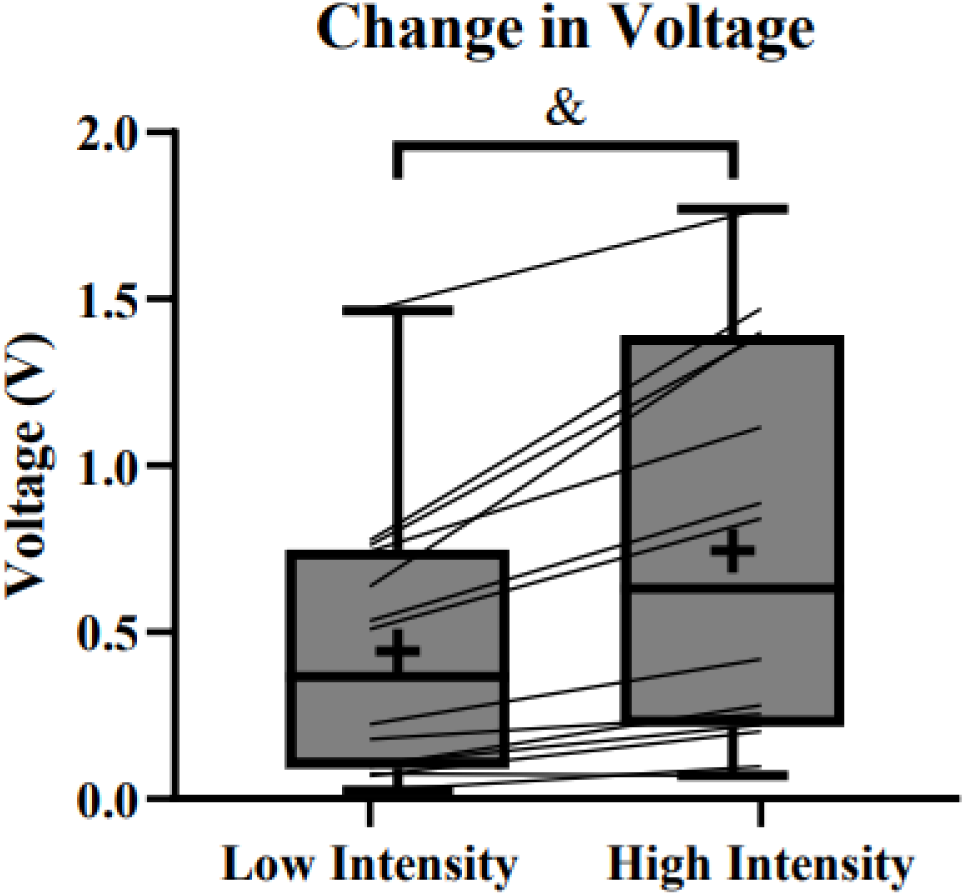
Voltage between intensities. The increase in voltage supplied to the kneecap was observed from low intensity to high intensity conditions. & indicates a significant difference.

### Average pedal power, torque, NSI and variability during cycling

#### Power at Higher-Intensity Cycling

A two-way repeated-measures ANOVA revealed a significant condition × leg interaction (p=0.017, ηp²=0.232). Follow-up comparisons demonstrated a significant reduction in power output in the pain leg during the EXP compared with the CNTRL condition (p=0.010, d=0.657), whereas no significant differences were observed in the non-pain leg (p=0.115, d=0.342) (Figure 3A). For this measure, NSI differed significantly between conditions (p=0.005, d=0.659), indicating greater reliance on the non-pain leg during EXP (Figure 3B). For CV between conditions, there was no significant effect of leg (p=0.523) or condition (p=0.755), nor was there any significant interaction (p=0.871).

**Figure 3.**
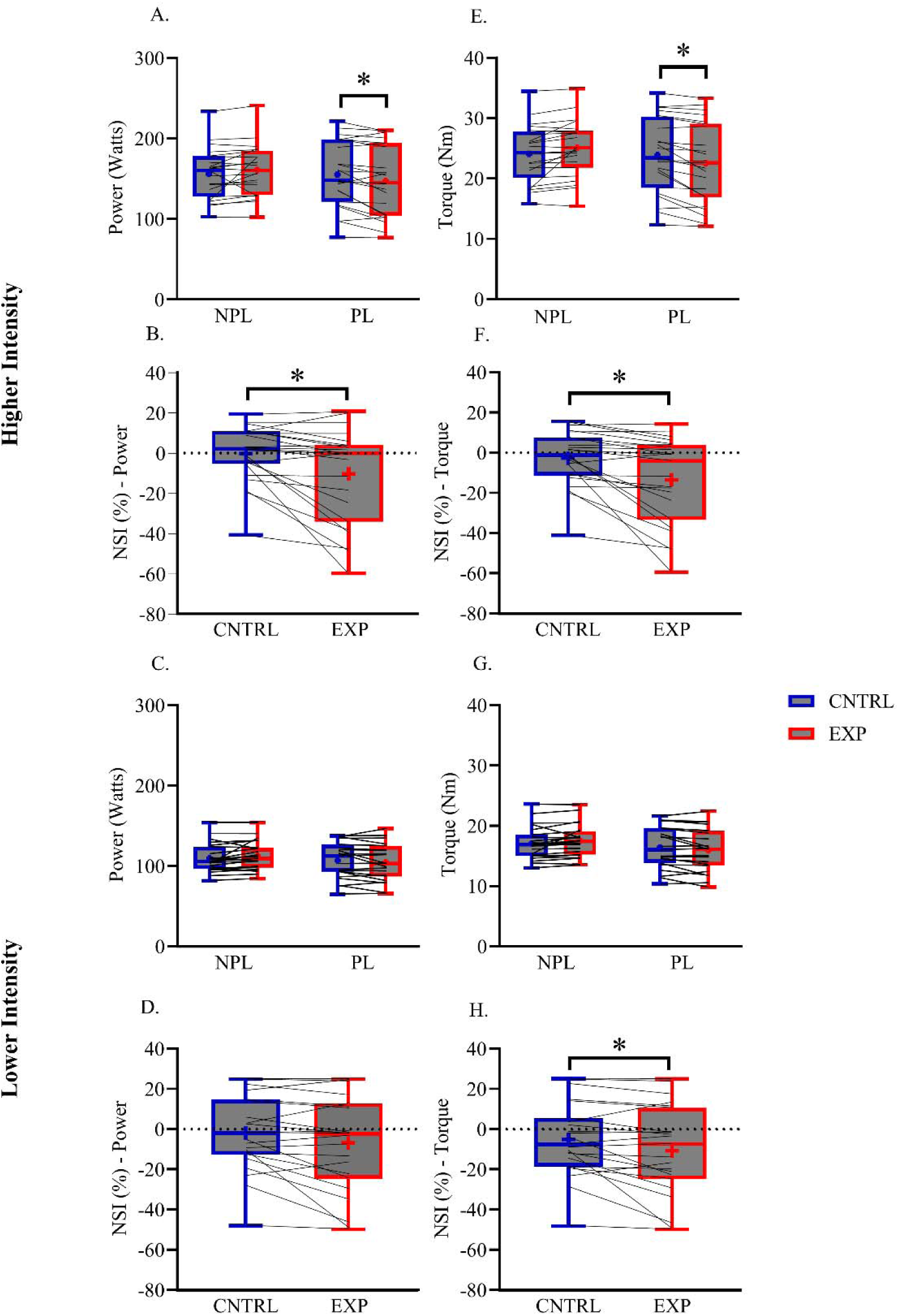
Effects of pain on power, torque and NSI. **(A)** HI power leg comparison between conditions; **(B)** HI NSI for power between conditions; **(C)** LI power leg comparison between conditions; **(D)** LI NSI for power between conditions; **(E)** HI torque leg comparison between conditions; **(F)** HI NSI for torque between conditions; **(G)** LI torque leg comparison between conditions; **(H)** LI NSI for torque between conditions; * indicates a significant difference between conditions; + indicates mean of the data; CNTRL: control condition; EXP: experimental condition; HI: higher intensity; LI: lower intensity; NSI: normalized symmetry index.

#### Power at Lower-Intensity Cycling

A significant condition × leg interaction was identified (p=0.036 ηp²=0.186); however, post hoc comparisons did not reveal significant within-leg differences for either the pain leg (p=0.154, d=0.387) or non-pain leg (p=0.093, d=0.367) (Figure 3C). For this measure, no significant difference in NSI was observed between conditions (p=0.083, d=0.449) (Figure 3D). For CV, there was no leg effect (p=0.634), condition effect (p=0.379), or interaction effect (p=0.357).

#### Torque at Higher-Intensity Cycling

A significant condition × leg interaction was observed for torque (p<0.001, ηp²=0.397). Post hoc analyses demonstrated a significant reduction in torque production in the pain leg during EXP conditions compared with the CNTRL condition (p=0.026, d=0.566), whereas no significant changes were observed in the non-pain leg (p=0.737, d=0.068) (Figure 3E). NSI differed significantly between conditions for (p<0.001, d=0.754), indicating increased reliance on the non-pain leg during the EXP condition (Figure 3F). For CV, there was no leg effect (p=0.280), condition effect (p=0.910), or interaction effect (p=0.963).

#### Torque at Lower-Intensity Cycling

A significant condition × leg interaction was present (p=0.017, ηp²=0.232); however, post hoc comparisons revealed no significant within-leg differences for either the pain leg (p=0.104, d=0.427) or non-pain leg (p=0.146, d=0.225) (Figure 3G). However, a significant NSI was observed for this measure (p=0.017, d=0.538), where NSI demonstrated greater reliance on the non-pain leg (Figure 3H). For CV, there was no leg effect (p=0.235), condition effect (p=0.488), or interaction effect (p=0.317).

### Handlebar and Seat Post

#### Higher-Intensity Cycling

Handlebar force showed no significant main effect of condition (p=0.519), side (p=0.482) or condition × side interaction (p=0.758). Seat post force also did not differ significantly between EXP and CNTRL conditions (p=0.519, d=0.137).

#### Lower-Intensity Cycling

For handlebar force the two-way repeated-measures ANOVA revealed no significant effect of condition (p=0.993), side (p=0.326) or condition × side interaction (p=0.449). Seat-post force similarly did not differ between EXP and CNTRL conditions (p=0.298, d=0.222). Mean handlebar and seat-post force values are reported in Table A.4 and A.5, respectively.

### Individual classifications for average power, torque, ΔNSI% and ΔCV%

#### During Higher-Intensity Cycling

For power, five participants demonstrated large reductions in pain-leg power [mean (SD), -19.1 (3.50) %], four demonstrated moderate reductions (–8.67 (3.40) %), ten demonstrated small reductions (–1.89 (1.20) %) and four demonstrated positive responses (5.23 (2.78) %) (Figure 4A and B). For torque, five participants demonstrated large reductions (−18.76 (2.43) %), four moderate reductions (−9.17 (3.04) %), eleven small reductions (−1.93 (1.13) %), and three increases (1.51 (0.59) %) (Figure 4A, D). ΔNSI showed large redistribution towards the non-pain leg in four participants (−38.6 (11.1) %), moderate redistribution towards in two (−19.6 (1.1) %), and small redistribution in eleven (−5.135 (3.327) %). Six participants instead demonstrated redistribution towards pain-leg (3.33 (2.73) %) (Figure 5A and B). For the same parameter, ΔCV%, four participants demonstrated large reductions in variability (−29.24 (4.11) %), nine demonstrated moderate reductions in variability (−11.55 (7.93) %), one demonstrated minimal variability change (0.24%), four demonstrated moderate increases in variability (18.27 (6.74) %), and five demonstrated large increases in variability (42.645 (15.11) %) (Figure 6A and B).

**Figure 4.**
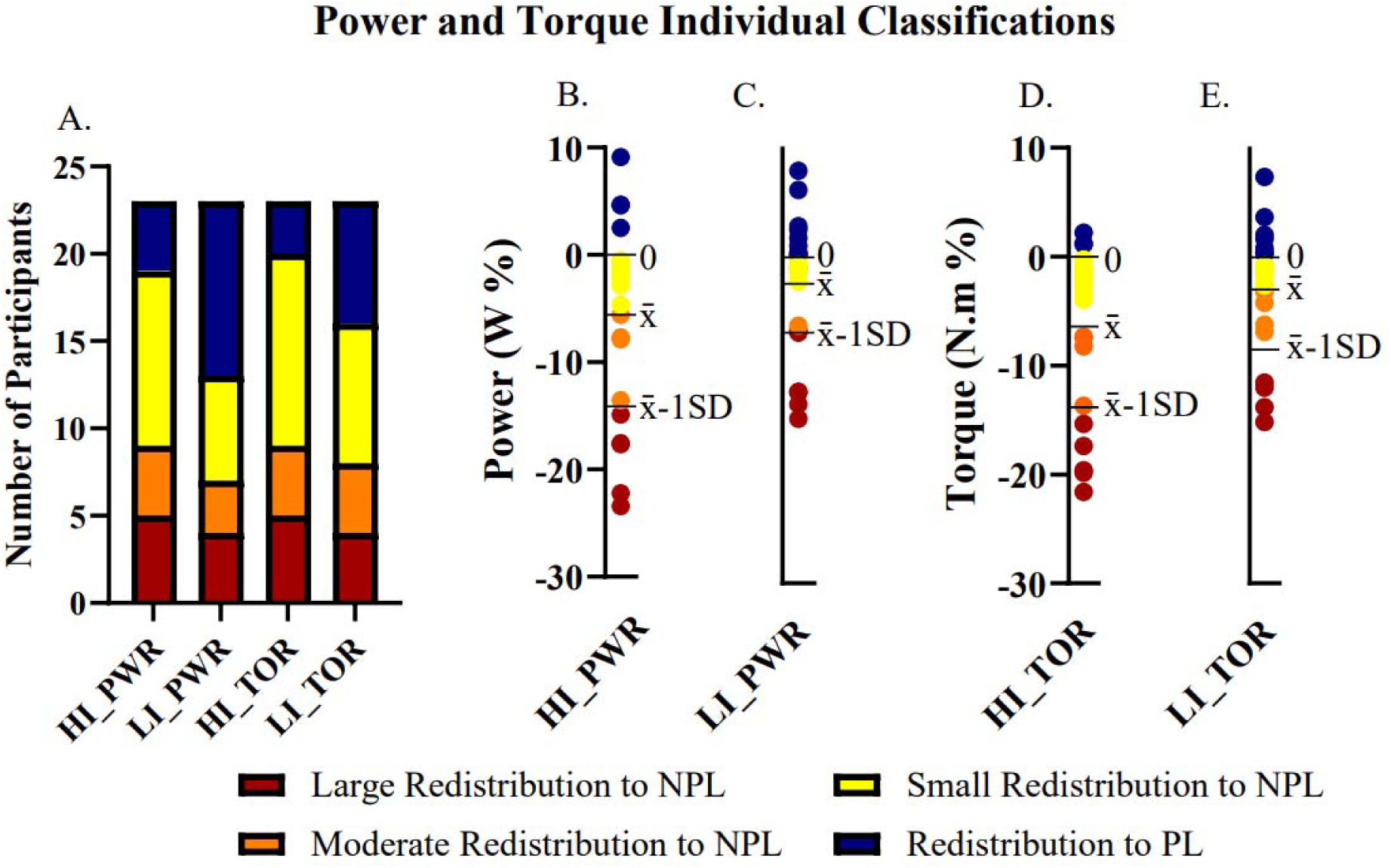
Individual response classification and participant data for power and torque. **(A)** Number of participants classified in each category for power and torque at higher and lower intensities. **(B)** Individual data points for HI_PWR. **(C)** Individual data points for LI_PWR. **(D)**Individual data points for participant responses for HI_TOR. **(E)** Ind –: mean.

**Figure 5.**
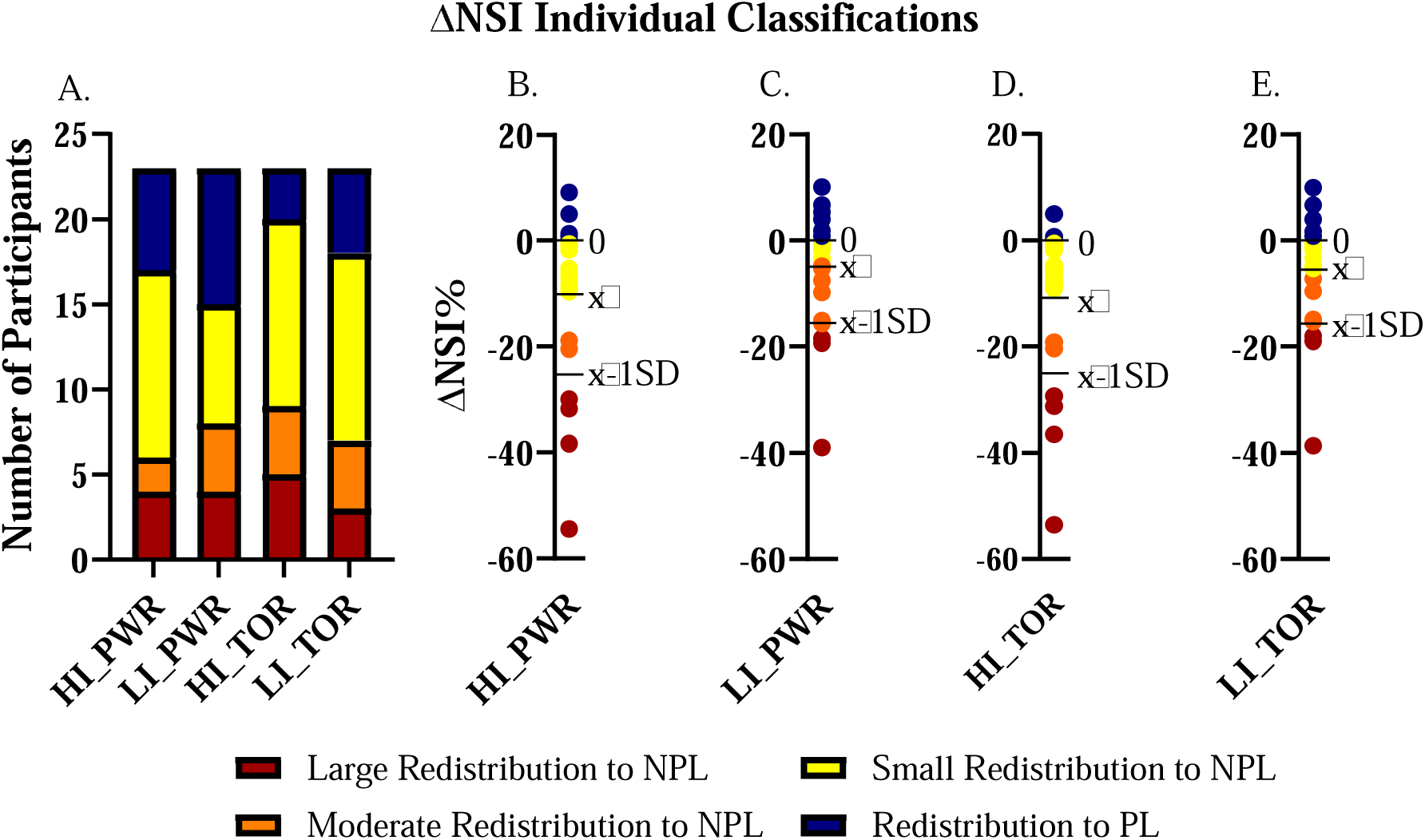
Individual response classification and participant data for power and torque ΔNSI. **(A)** Number of participants classified in each category for power and torque ΔNSI at higher and lower intensities. **(B)** Individual data points for HI_PWR. **(C)** Individual data points for LI_PWR. **(D)** Individual data points for participant responses for HI_TOR. **(E)** Individual data points for LI_TOR. HI_TOR: higher intensity torque; HI_PWR: higher intensity power; LI_PWR: lower intensity power; LI_TOR: lower intensity torque; NPL: non-pain leg; PL: pain leg; x□: mean.

**Figure 6.**
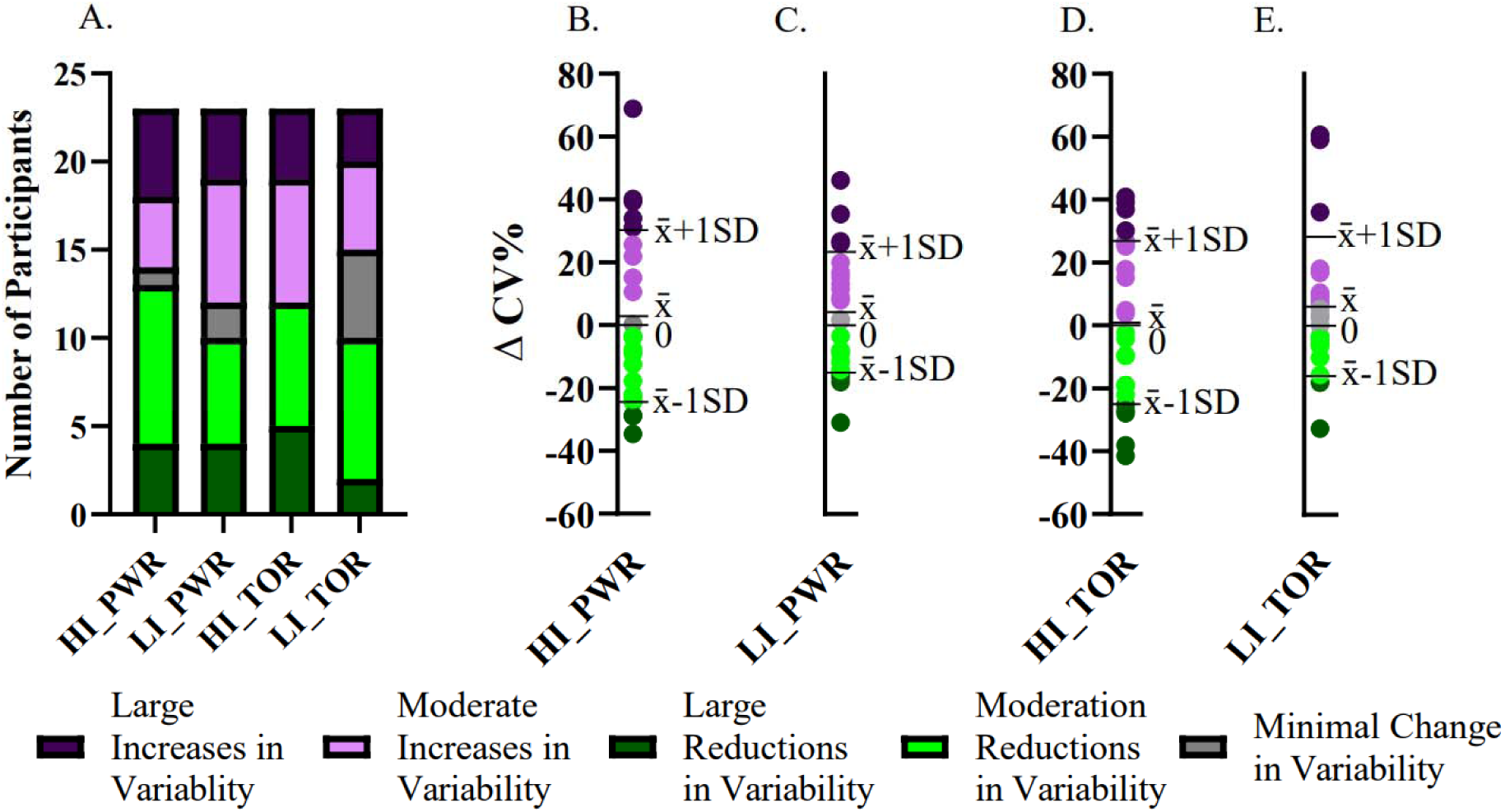
Individual response classification and participant data for power and torque ΔCV%. **(A)** Number of participants classified in each category for power and torque ΔCV% at higher and lower intensities. **(B)** Individual data points for HI_PWR. **(C)** Individual data points for LI_PWR. **(D)** Individual data points for participant responses for HI_TOR. **(E)** Individual data points for LI_TOR. HI_TOR: higher intensity torque; HI_PWR: higher intensity power; LI_PWR: lower intensity power; LI_TOR: lower intens –: mean.

For torque, five participants had large negative responses (−18.76 (2.43) %), four participants had moderate negative responses (−9.17 (3.04) %), eleven demonstrated small negative responses (–1.93 (1.13) %) and three participants demonstrated positive responses (1.51 (0.59) %) (Figure 4A and D). For ΔNSI, four participants demonstrated large redistribution towards the non-pain leg (−37.7 (11.04) %), two demonstrated moderate redistribution toward the non-pain leg (−19.75 (0.85) %), thirteen demonstrated small redistribution toward the non-pain leg (−5.14 (3.18) %), and four demonstrated redistribution toward the pain leg (1.62 (2.22) %) (Figure 5A and D). For the same parameter, ΔCV%, five participants demonstrated large reductions in variability (−32.0 (7.30) %), seven demonstrated moderate reductions in variability (−12.34 (7.78) %), seven demonstrated moderate increases in variability (17.080 (9.753) %), and four demonstrated large increases in variability (36.8 (4.68) %). No participants demonstrated minimal variability change (Figure 6A and D).

#### During Lower-Intensity Cycling

For power, four participants demonstrated large negative responses (mean (SD), -13.3 (1.10) %), 3 as moderate negative responses (–6.62 (0.31) %), 6 as small negative responses (–1.07 (0.66) %) and ten as positive responses (2.37 (2.70) %) (Figure 4A and C). For ΔNSI, four participants demonstrated large redistribution towards the non-pain leg (−23.1 (10.7) %), four demonstrated moderate redistribution towards the non-pain leg (−9.39 (4.2) %), seven demonstrated small redistribution towards the non-pain leg (−2.384 (1.570) %), and eight demonstrated redistribution towards pain-leg (4.26 (3.13) %) (Figure 5A and C). For power ΔCV%, four participants demonstrated large reductions in variability (−20.2 (7.2) %), six demonstrated moderate reductions in variability (−9.08 (3.67) %), two demonstrated minimal variability change (1.76 (0.22) %), four demonstrated large increases in variability (33.6 (9.30) %), and seven demonstrated moderate increases in variability (13.4 (4.30) %) (Figure 6A and C). For torque, four participants had large negative responses (−13.1 (1.68) %), four participants had moderate negative responses (−5.10 (1.72) %), eight demonstrated small negative responses (–1.47 (0.71) %) and seven participants demonstrated positive responses (2.37 (2.75) %) (Figure 4A and E). For ΔNSI, three participants demonstrated large redistribution towards the non-pain leg (−25.2 (11.7) %), four demonstrated moderate redistribution toward the non-pain leg (−11.7 (4.0) %), eleven demonstrated small redistribution toward the non-pain leg (−2.55 (1.92) %), and five demonstrated redistribution toward the pain leg (4.60 (3.74) %) (Figure 5A and E). For torque ΔCV%, two participants demonstrated large reductions in variability (−25.3 (10.3) %), eight demonstrated moderate reductions in variability (−7.00 (4.03) %), five demonstrated moderate increases in variability (12.6 (5.06) %), and three demonstrated large increases in variability (52.0 (13.8) %). Five participants demonstrated minimal variability change (5.68 (6.55) %) (Figure 6A and E).

## Discussion

This study investigated the effects of movement-evoked experimental knee pain on bilateral cycling biomechanics and characterized inter-individual variability in these responses. Overall, group-level analyses indicated that pain reduced power and torque production in the pain leg and shifted inter-limb symmetry toward the non-pain leg, with the clearest effects observed at higher intensities. At lower intensity, pain-leg power and torque did not differ significantly between EXP and CNTRL conditions; however, torque NSI shifted toward the non-pain leg, suggesting that bilateral redistribution may occur even in the absence of statistically significant changes within either limb individually. However, substantial inter-individual variability was observed, ranging from large reductions in pain leg power and torque to minimal changes or even increased reliance on the pain leg. Together, these findings indicate that movement-evoked knee pain can alter bilateral mechanical output during cycling, but that the resulting motor adaptation is highly individualized.

At the group level, the reduction in pain-leg torque and power is broadly consistent, with more pronounced effects at higher intensities. The observed pattern is consistent with inhibitory motor strategies that are proposed in pain adaptation models (Merkle, Sluka, & Frey-Law, 2018), suggesting that pain may modify motor output in a manner that reduces mechanical demand on the painful region while allowing the task to continue (Hodges & Tucker, 2011; Lund, Donga, Widmer, & Stohler, 1991). Within the context of cycling, this was reflected in a reduced mechanical contribution from the pain leg and a relative shift toward the contralateral leg. This interpretation is corroborated by the strength inhibition theory, which proposes that individuals reduce force-generating capacity in response to pain, resulting in decreased mechanical output during task execution (Merkle, Sluka, & Frey-Law, 2018; Lund, Donga, Widmer, & Stohler, 1991). These results are also consistent with the results from Charlton et al. (Charlton, et al., 2024) and Cabral et al. (Cabral, et al., 2023), which indicated a reduction in joint movement and force production, respectively, under pain (Charlton, et al., 2024; Cabral, et al., 2023). These findings extend this previous work by demonstrating that knee pain can also modify bilateral loading during a dynamic, repetitive, multi-joint task such as cycling. However, because muscle activation and other neurophysiological measures were not assessed, the mechanisms underlying the observed reductions in mechanical output cannot be determined from the present study.

The effects were most evident during the higher-intensity/higher-pain condition, in which pain-leg power and torque were both significantly reduced and NSI shifted toward the non-pain leg. These findings should not, however, be interpreted as evidence of a workload-dependent effect alone. In the present experimental design, cycling workload and target pain intensity increased concurrently: the lower-intensity condition was performed at 20% PPO with pain targeted at 1–2/10, whereas the higher-intensity condition corresponded to a greater workload with pain targeted at 5–6/10. Consequently, the larger biomechanical changes observed in the higher condition may reflect the greater pain intensity, greater mechanical demand, or an interaction between the two. Future studies that independently manipulate workload and pain intensity are required to determine their respective contributions to pain-related cycling adaptations.

The NSI findings provide additional insight into bilateral compensation. The present study demonstrated that experimental knee pain altered NSI toward the non-pain leg during higher-intensity/higher-pain cycling. Redistribution toward the contralateral limb has also been reported in individuals with knee osteoarthritis and is commonly interpreted as a strategy to reduce loading of the affected limb (Liu, Amiri, McGregor, & Bull, 2022). Our present findings suggest that comparable compensatory strategies can occur acutely in healthy individuals in response to experimentally induced pain. Of particular interest, torque NSI was significantly altered during the lower-intensity condition despite the absence of significant within-leg differences in torque. This suggests that symmetry-based measures may capture relatively small, coordinated changes across both limbs that may not be apparent when each limb is considered independently. Accordingly, bilateral metrics such as NSI may provide complementary information when evaluating pain-related changes in cycling mechanics. These results are supported by Hodges & Tucker (Hodges & Tucker, 2011), who suggest that when exposed to pain, the nervous system prioritizes early protective adaptations prior to large performance decrements. Thus, NSI may provide additional information about pain-related changes in motor output that may be overlooked when examining limb-specific measures alone.

In contrast to the lower-limb findings, no significant differences were observed between EXP and CNTRL conditions for handlebar or seat-post mechanical output variables. This suggests that the most consistent group-level adaptations were expressed through changes in lower-limb mechanical output rather than measurable redistribution through the upper body. However, the absence of significant differences in handlebar or seat-post force should not be interpreted as evidence that upper body compensation did not occur. Changes in trunk posture, muscle activation, coordination or force direction may not have been captured by the selected measures. Additionally, heterogeneous compensatory strategies between individuals could reduce the likelihood of detecting consistent group-level effects.

A key finding of this study was the substantial inter-individual variability in biomechanical responses to pain. While group-level response indicated a reduced mechanical contribution from the pain leg, this pattern was not universal. During the higher-intensity condition, most participants reduced pain-leg power and torque, whereas four participants increased pain-leg power and three increased pain-leg torque. This is consistent with prior investigations reporting marked variability in motor responses to experimentally induced knee pain during locomotion (Charlton, et al., 2024) (Dionne, Bartoshuk, Mogil, & Witter, 2005). Rather than eliciting a single stereotypical response, pain appears to permit multiple motor solutions through which individuals balance task performance and protection from perceived threat (Merkle, Sluka, & Frey-Law, 2018) (Hodges & Tucker, 2011).

This heterogeneity also highlights an important limitation of relying exclusively on group-level analyses. A statistically significant mean reduction in pain-leg output may accurately describe the predominant response while obscuring individuals who demonstrate little change or adapt in the opposite direction. At the same time, individual classifications should be interpreted cautiously. As emphasized by Atkinson et al. (Atkinson, Williamson, & Batterham, 2019; Dionne, Bartoshuk, Mogil, & Witter, 2005), categorizing individuals as “responders” or “non-responders” may oversimplify continuous biological variability. Indeed, findings from the present study demonstrate a continuum of motor response strategies in response to painful stimulus within the same cohort, that might not be detected through group-level analyses.

Individual changes in movement variability further support this interpretation. Although no significant group-level changes in movement variability were observed, individual responses to movement-evoked pain showed both substantial increases and reductions in torque and power variability. These opposing responses may represent different approaches to movement regulation under painful conditions. Reduced variability might reflect a constrained motor strategy aimed at stabilizing movement and minimizing the threat of pain, whereas increased variability may reflect exploratory behavior to identify less painful movement solutions (Merkle, Sluka, & Frey-Law, 2018). Overall, these variabilities might reflect differences in pain sensitivity, nociceptive processing, fear-related behaviors, and individual motor response strategies (Aboushaar & Serrano, 2024). Because these mechanisms were not directly assessed, such interpretations remain speculative and should be investigated in future studies.

There are several methodological considerations with the current study that should be acknowledged. Pain intensity reported by participants was self-reported, which may have introduced variability in nociceptive exposure. In addition, classification thresholds used in the current study were derived relative to the sampled distribution rather than established clinical cutoffs; therefore, these results cannot be interpreted as definitive physiological categories. Consistent with recommendations from Atkinson et al. (Atkinson, Williamson, & Batterham, 2019), these classifications were intended to describe the spectrum of observed responses rather than identifying distinct responder phenotypes. Although movement-evoked pain using electrical stimulation has proved successful for mimicking knee pain in previous studies (Cabral, et al., 2023), it does not fully replicate the total complexity of real clinical conditions, including chronic neuromuscular adaptations, inflammation and psychosocial factors. Furthermore, because the CNTRL condition involved no electrical stimulation, sensory, attentional, or anticipatory effects associated with electrical stimulation could not be completely separated from pain-specific effects. The study included both females and males but was not designed or sufficiently powered to examine sex-related differences, which may contribute to the observed inter-individual variability in pain responses. Lastly, it is important to recognize that the present study quantified behavioral motor responses to pain through external biomechanical measures, including force, torque, power, limb asymmetry and movement variability. Although these behavioral responses likely reflect modifications in neuromuscular control, no direct neurophysiological measures were obtained. Consequently, the underlying neurophysiological mechanisms responsible for these responses cannot be determined from the present data. Future studies combining bilateral biomechanical measurements with electromyographic and neurophysiological assessments would help identify the mechanisms underlying different individual motor strategies.

## Conclusion

The findings of this study demonstrate that movement-evoked knee pain alters cycling biomechanics, particularly at higher intensities, by reducing power and torque contribution from the painful limb while increasing reliance on the non-painful limb. In addition to these group-level effects, substantial inter-individual differences existed in magnitude, direction and variability of movement responses. Some participants demonstrated clear unloading and redistribution strategies while others maintained relatively symmetrical output despite exposure to pain. These findings support current theories suggesting that pain does not elicit a single universal motor response, but rather a broad spectrum of movement responses aimed at balancing task performance with protection from pain. Collectively, these results highlight the importance of incorporating individual-level analyses when examining pain-related motor behavior and may have important implications for developing individualized rehabilitation approaches that rely on cycling-based exercise. Importantly, the present findings demonstrate the value of evaluating bilateral mechanical output and individual response patterns when investigating pain-related movement adaptations. This may be particularly relevant to clinical populations such as individuals with knee osteoarthritis, in whom persistent inter-limb asymmetry and preferential loading of the non-painful limb may contribute to altered joint loading and movement strategies. By demonstrating that acute experimental knee pain can produce similar redistribution of lower-limb mechanical output during dynamic cycling, the present study provides a framework for investigating how pain-related asymmetry emerges and varies between individuals. This study has important implications for rehabilitation and exercise prescription for asymmetry associated with pain. These findings may also inform future approaches to cycling-based rehabilitation by highlighting the importance of monitoring both limbs and considering individual responses when prescribing and evaluating exercise in the presence of pain.

## Abbreviations

Abbreviation Definition

NSI: Normalized Symmetry Index
DAQ: Data Acquisition
ICC: Intraclass Correlation Coefficient
SD: Standard Deviation
PPO: Peak Power Output
W: Watts
RPM: Revolutions Per Minute
EXP: Experimental Condition
CNTRL: Control Condition
LI_CNTRL: Lower-Intensity Control
LI_EXP: Lower-Intensity Experimental
HI_CNTRL: Higher-Intensity Control
HI_EXP: Higher-Intensity Experimental
NPL: Non-Pain Leg
PL: Pain Leg
EXP_PS: Experimental Pain Side
EXP_NPS: Experimental Non-Pain Side
CNTRL_PS: Control Pain Side
CNTRL_NPS: Control Non-Pain Side
CV: Coefficient of Variation
LI: Lower Intensity
HI: Higher Intensity
N: Newton
N.m: Newton Meter

## Supplementary Material

**Table A.1.**
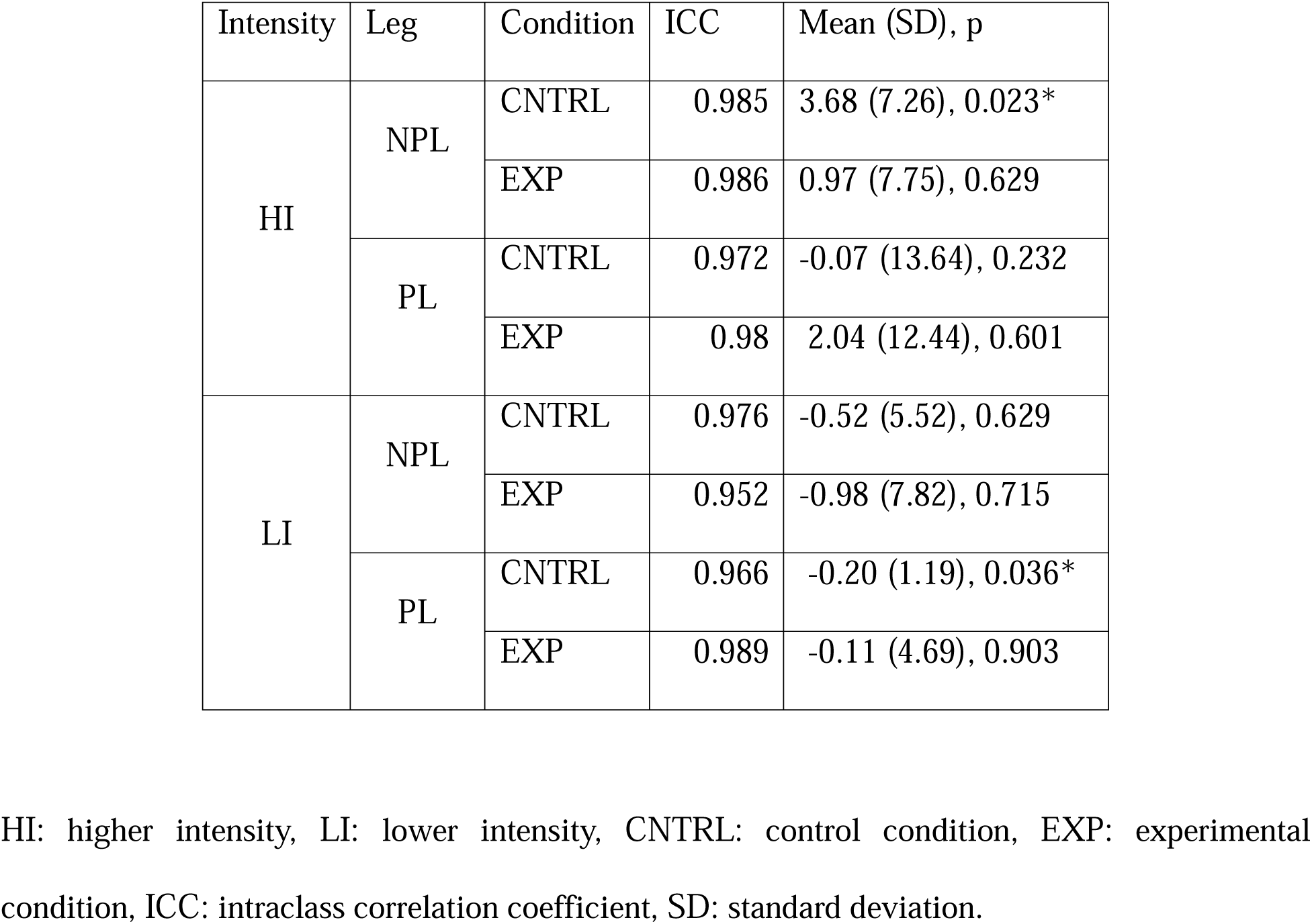
Reliability Results for Power.

| Intensity | Leg | Condition | ICC | Mean (SD), p |
| --- | --- | --- | --- | --- |
| HI | NPL | CNTRL | 0.985 | 3.68 (7.26), 0.023* |
|  |  | EXP | 0.986 | 0.97 (7.75), 0.629 |
|  | PL | CNTRL | 0.972 | -0.07 (13.64), 0.232 |
|  |  | EXP | 0.98 | 2.04 (12.44), 0.601 |
| LI | NPL | CNTRL | 0.976 | -0.52 (5.52), 0.629 |
|  |  | EXP | 0.952 | -0.98 (7.82), 0.715 |
|  | PL | CNTRL | 0.966 | -0.20 (1.19), 0.036* |
|  |  | EXP | 0.989 | -0.11 (4.69), 0.903 |

HI: higher intensity, LI: lower intensity, CNTRL: control condition, EXP: experimental condition, ICC: intraclass correlation coefficient, SD: standard deviation.

**Table A.2.**
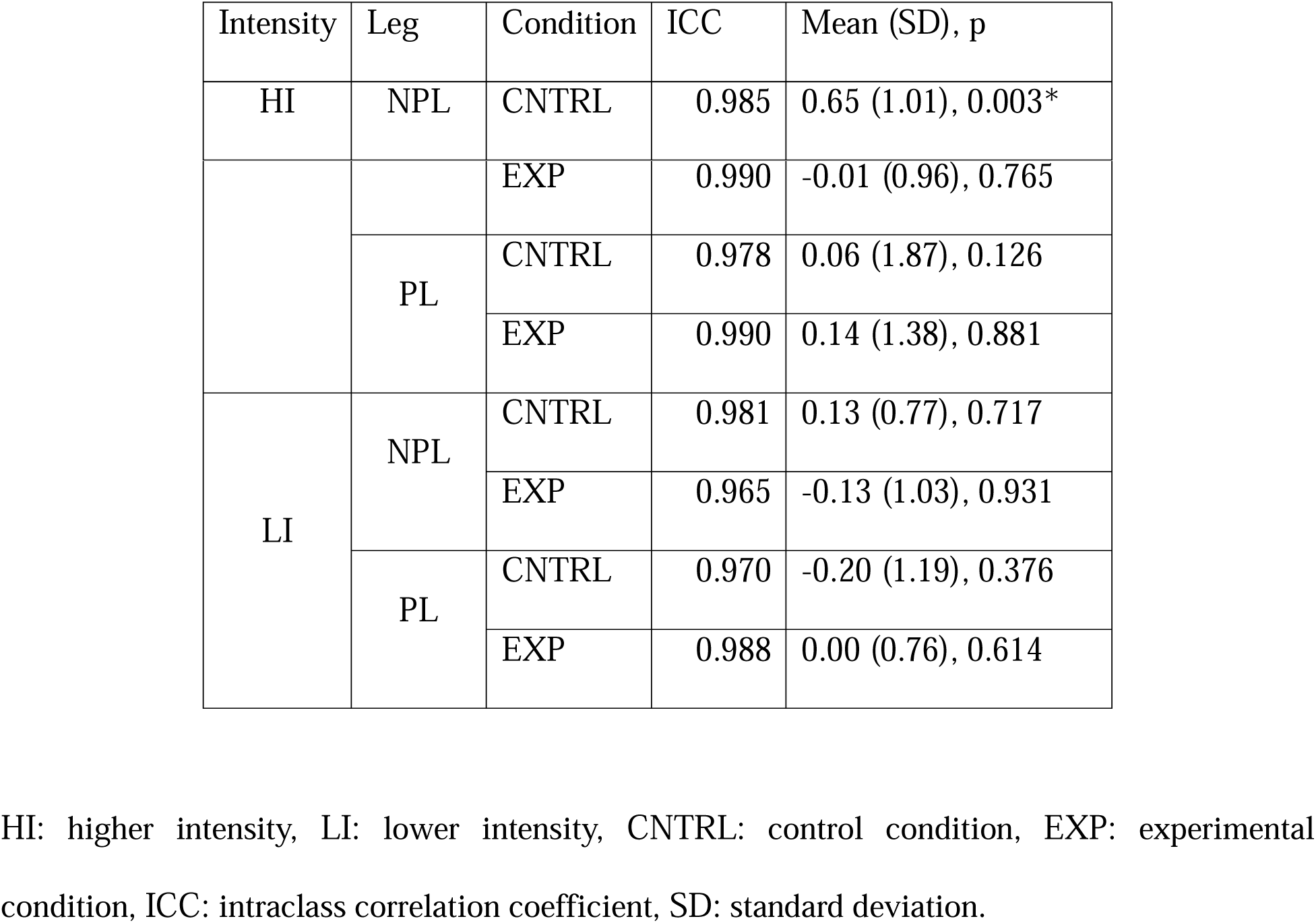
Reliability Results for Torque.

**Table A.3.**
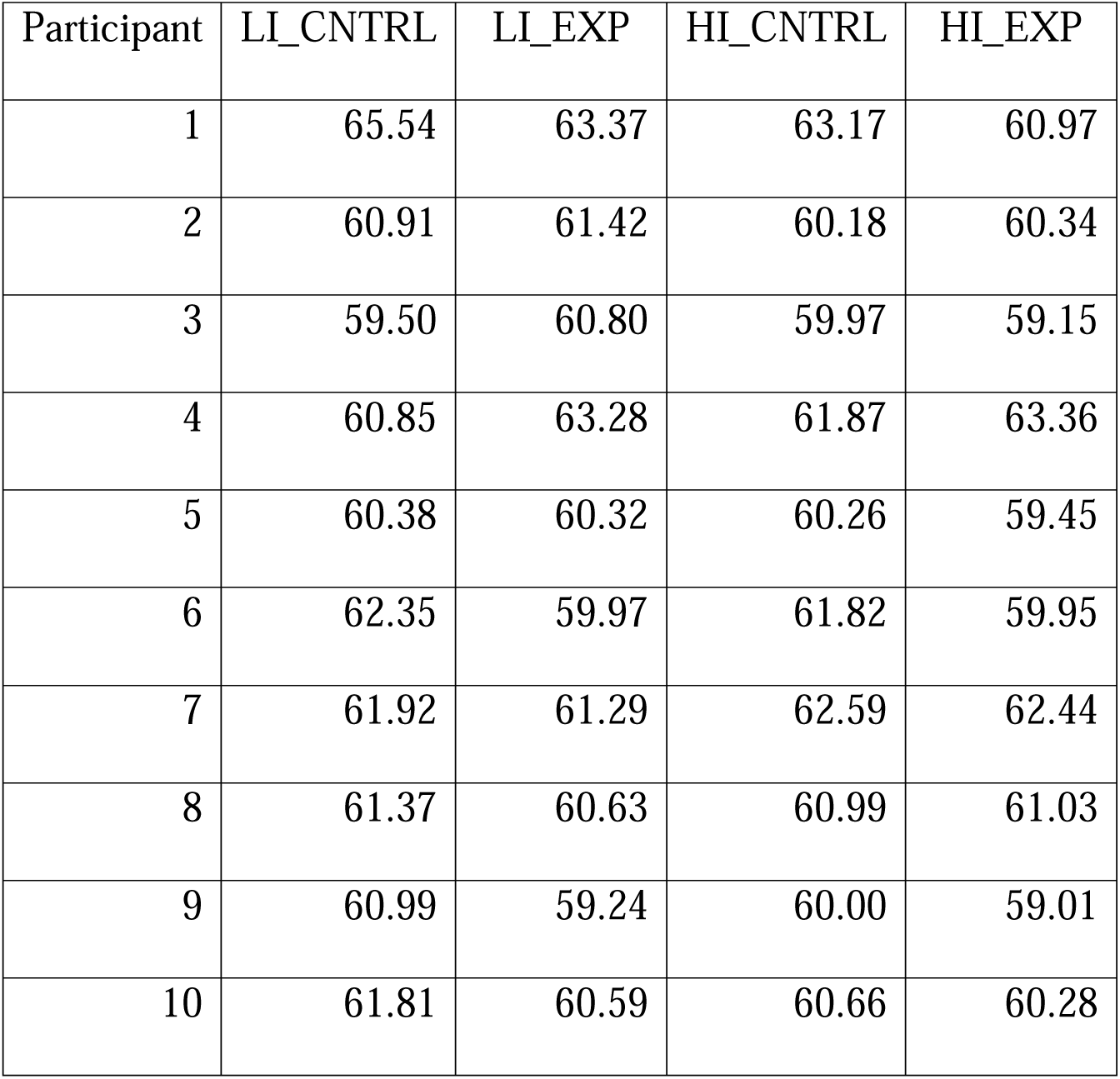
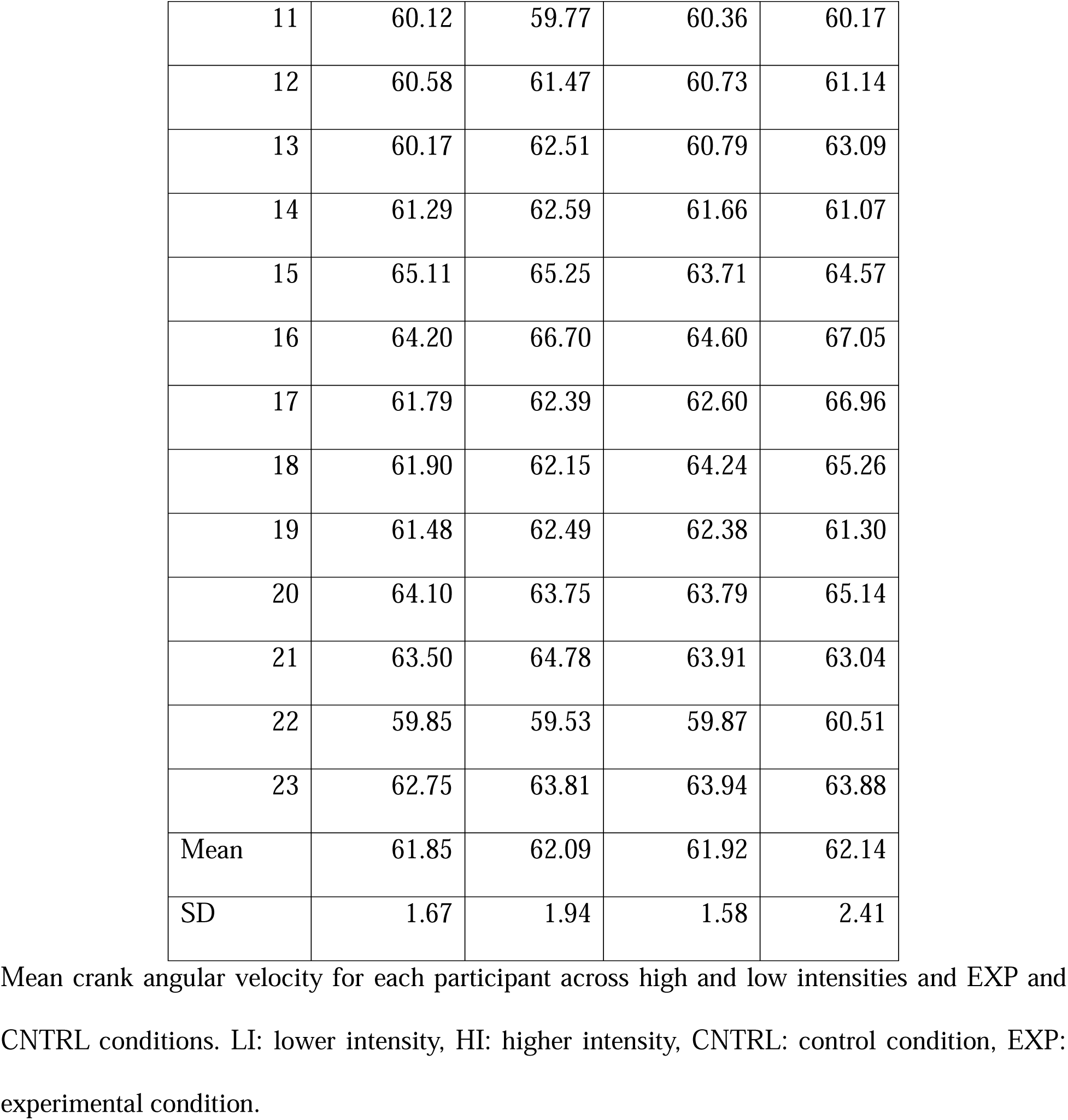
Mean Crank Angular Velocity.

| Participant | LI_CNTRL | LI_EXP | HI_CNTRL | HI_EXP |
| --- | --- | --- | --- | --- |
| 1 | 65.54 | 63.37 | 63.17 | 60.97 |
| 2 | 60.91 | 61.42 | 60.18 | 60.34 |
| 3 | 59.50 | 60.80 | 59.97 | 59.15 |
| 4 | 60.85 | 63.28 | 61.87 | 63.36 |
| 5 | 60.38 | 60.32 | 60.26 | 59.45 |
| 6 | 62.35 | 59.97 | 61.82 | 59.95 |
| 7 | 61.92 | 61.29 | 62.59 | 62.44 |
| 8 | 61.37 | 60.63 | 60.99 | 61.03 |
| 9 | 60.99 | 59.24 | 60.00 | 59.01 |
| 10 | 61.81 | 60.59 | 60.66 | 60.28 |
| 11 | 60.12 | 59.77 | 60.36 | 60.17 |
| 12 | 60.58 | 61.47 | 60.73 | 61.14 |
| 13 | 60.17 | 62.51 | 60.79 | 63.09 |
| 14 | 61.29 | 62.59 | 61.66 | 61.07 |
| 15 | 65.11 | 65.25 | 63.71 | 64.57 |
| 16 | 64.20 | 66.70 | 64.60 | 67.05 |
| 17 | 61.79 | 62.39 | 62.60 | 66.96 |
| 18 | 61.90 | 62.15 | 64.24 | 65.26 |
| 19 | 61.48 | 62.49 | 62.38 | 61.30 |
| 20 | 64.10 | 63.75 | 63.79 | 65.14 |
| 21 | 63.50 | 64.78 | 63.91 | 63.04 |
| 22 | 59.85 | 59.53 | 59.87 | 60.51 |
| 23 | 62.75 | 63.81 | 63.94 | 63.88 |
| Mean | 61.85 | 62.09 | 61.92 | 62.14 |
| SD | 1.67 | 1.94 | 1.58 | 2.41 |

Mean crank angular velocity for each participant across high and low intensities and EXP and CNTRL conditions. LI: lower intensity, HI: higher intensity, CNTRL: control condition, EXP: experimental condition.

**Table A.4.** Mean Force Values for Handlebar Data (N)

| Condition | Higher Intensity<br>(Mean (SD)) | Lower Intensity<br>(Mean (SD)) |
| --- | --- | --- |
| EXP_PS | 46.77 (18.16) | 51.50 (19.78) |
| EXP_NPS | 45.39 (16.63) | 48.99 (19.92) |
| CNTRL_PS | 45.89 (14.21) | 51.03 (18.86) |
| CNTRL_NPS | 45.03 (17.13) | 49.44 (20.85) |

Note: EXP\_PS: experimental condition pain side, EXP\_NPS: experimental condition non-pain side, CNTRL\_PS: control condition pain side, CNTRL\_NPS: control condition non-pain side.

**Table A.5.** Mean Values for Seat Post Mediolateral Sway Data (N)

| Condition | Value (Mean (SD)) |
| --- | --- |
| LI_EXP | -2.16 (26.20) |
| LI_CNTRL | -0.927 (24.10) |
| HI_EXP | 1.25 (26.10) |
| HI_CNTRL | -0.836 (28.10) |

Note: LI\_CNTRL: lower-intensity control condition, LI\_EXP: lower-intensity experimental condition, HI\_EXP: higher-intensity experimental condition, HI\_CNTRL: higher-intensity control condition.

